# Latent motives guide structure learning during adaptive social choice

**DOI:** 10.1101/2020.06.06.137893

**Authors:** Jeroen M. van Baar, Matthew R. Nassar, Wenning Deng, Oriel FeldmanHall

**Author notes:** Corresponding author at 349 Metcalf Hall, 190 Thayer St, Providence, RI 02912, United States, **Email**. **Author Contributions** J.v.B. and O.F.H. designed research. J.v.B. and W.D. performed research. J.v.B., M.N., and O.F.H. analyzed data. J.v.B. and O.F.H. wrote the paper. J.v.B., M.N., and O.F.H. edited the manuscript.

## Abstract

Predicting the behavior of others is an essential part of human cognition that enables strategic social behavior (e.g., cooperation), and is impaired in multiple clinical populations. Despite its ubiquity, social prediction poses a generalization problem that remains poorly understood: We can neither assume that others will simply repeat their past behavior in new settings, nor that their future actions are entirely unrelated to the past. Here we demonstrate that humans solve this challenge using a structure learning mechanism that uncovers other people’s latent, unobservable motives, such as greed and risk aversion. In three studies, participants were tasked with predicting the decisions of another player in multiple unique economic games such as the Prisoner’s Dilemma. Participants achieved accurate social prediction by learning the hidden motivational structure underlying the player’s actions to cooperate or defect (e.g., that greed led to defecting in some cases but cooperation in others). This motive-based abstraction enabled participants to attend to information diagnostic of the player’s next move and disregard irrelevant contextual cues. Moreover, participants who successfully learned another’s motives were more strategic in a subsequent competitive interaction with that player, reflecting that accurate social structure learning can lead to more optimal social behaviors. These findings demonstrate that advantageous social behavior hinges on parsimonious and generalizable mental models that leverage others’ latent intentions.

**Significance statement:** A hallmark of human cognition is being able to predict the behavior of others. How do we achieve social prediction given that we routinely encounter others in a dizzying array of social situations? We find people achieve accurate social prediction by inferring another’s hidden motives—motives that do not necessarily have a one-to-one correspondence with observable behaviors. Participants were able to infer another’s motives using a structure learning mechanism that enabled generalization. Individuals used what they learned about others in one setting to predict their actions in an entirely new setting. This cognitive process can explain a wealth of social behaviors, ranging from strategic economic decisions to stereotyping and racial bias.

## Introduction

Humans spend a significant amount of time trying to predict how others will behave. Which millennial has not agonized over the perfect emoji to round off a text message, hoping to elicit the desired response from a love interest? In a professional context, predicting how a co-worker will respond to feedback and adjusting a criticism accordingly can safeguard future collaborations. And even in large-scale societal coordination problems like political activism^1^, climate change mitigation^2^, and disease control^3^, reliable knowledge about the future intentions of others can be critical to reach the desired outcome.

Despite the ubiquity of social predictions in daily life, little is known about how humans solve this difficult task. Consider trying to predict whether a co-worker will be a good collaborator on a project. We cannot assume that our co-worker will act just as they did in the past, because each project is different (the people involved, the current economic conditions, etc.). Instead, we must selectively generalize what we know about our co-worker to this new situation. If we generalize too little from past experience, we deny ourselves potentially valuable social information, but generalizing too much makes us slow to respond to the changing behavior of others. To solve this generalization dilemma during social prediction, humans likely build parsimonious mental models of others’ behavior that are easy to update yet provide relatively accurate predictions in diverse social settings^4^. How do we construct and use these mental models to predict the intentions of others?

Here, we examine the possibility that motives, such as greed^5^, inequity aversion^6,7^, and risk aversion^8,9^, form the building blocks of these mental models of others’ behavior. Motives are plausible components of such models because they can predict behavior across contexts^10^, differ reliably between people^11–15^, and are inferred from an early age^16–18^. Studies of action understanding suggest that people can infer another’s motives by observing their actions^19^. For example, another’s food preference can be estimated by using Bayesian inference to compute which of several alternative foods is most likely to be selected^20^. However, this process becomes particularly challenging in naturalistic social settings because one behavior can arise from several motives^12,21^; for example, cooperation can stem from fairness concerns^6^, reputation management^22^, or warm-glow altruism^23^. To solve this inference problem, people must detect the latent structure of another’s choice patterns—i.e. their unobservable overarching goals and motives—across diverse social contexts.

Because it is unknown which cognitive mechanism supports learning of others’ latent motives, we test various computational accounts of this structure learning process, which can identify mechanisms that are not readily seen by examining behavior alone. Specifically, we evaluate whether a feature-based reinforcement learning system, thought to operate during non-social, low-dimensional structure learning tasks^24,25^, can support structural learning during social prediction. Rather than learn the value of observable features of the task (as is seen in non-social statistical structure learning work^24,26–30^), feature-based reinforcement learning requires participants to learn latent, unobservable features that can account for a person’s behavior, such as a motive like greed or risk aversion. If successful, latent structure learning would allow for greater efficiency in social prediction by selectively attending^24,26,31,32^ to task information that is specifically relevant to the inferred motive.

In our experimental Social Prediction Game, participants are tasked with predicting the choices of another Player interacting with anonymous Opponents (Fig. 1A) in four types of economic games, each characterized by distinct tensions between potential gains and losses: the Harmony Game (HG), Snowdrift Game (SG), Stag Hunt (SH), and Prisoner’s Dilemma (PD) (Fig. 1B). As Players move between games, the payoffs associated with cooperating and defecting shift. This causes the tensions associated with different underlying motives to vary from game to game, revealing several distinct and structured patterns of behavior^14^. For example, if a Player is motivated by risk aversion, they will choose to cooperate in a Snowdrift Game but defect in a Stag Hunt. Although these decisions appear contradictory at first blush, they are consistent at a latent motive level, as these actions both forgo the highest possible payoff in favor of minimizing potential losses^14,33^. Other motives, such as greed or envy, yield differently structured choice patterns across games^14^. This hidden structure allows us to test whether—and how—participants selectively generalize knowledge about others’ behavior to new social settings. We hypothesized that 1) people generalize information about others’ decisions between economic games on the basis of latent motives, 2) generalization is implemented by a feature-based reinforcement learning system that supports an increasing focusing of attention on information relevant to the inferred motives, and 3) people use these inferred motives to make more adaptive choices in subsequent competitive interactions.

**Figure 1.**
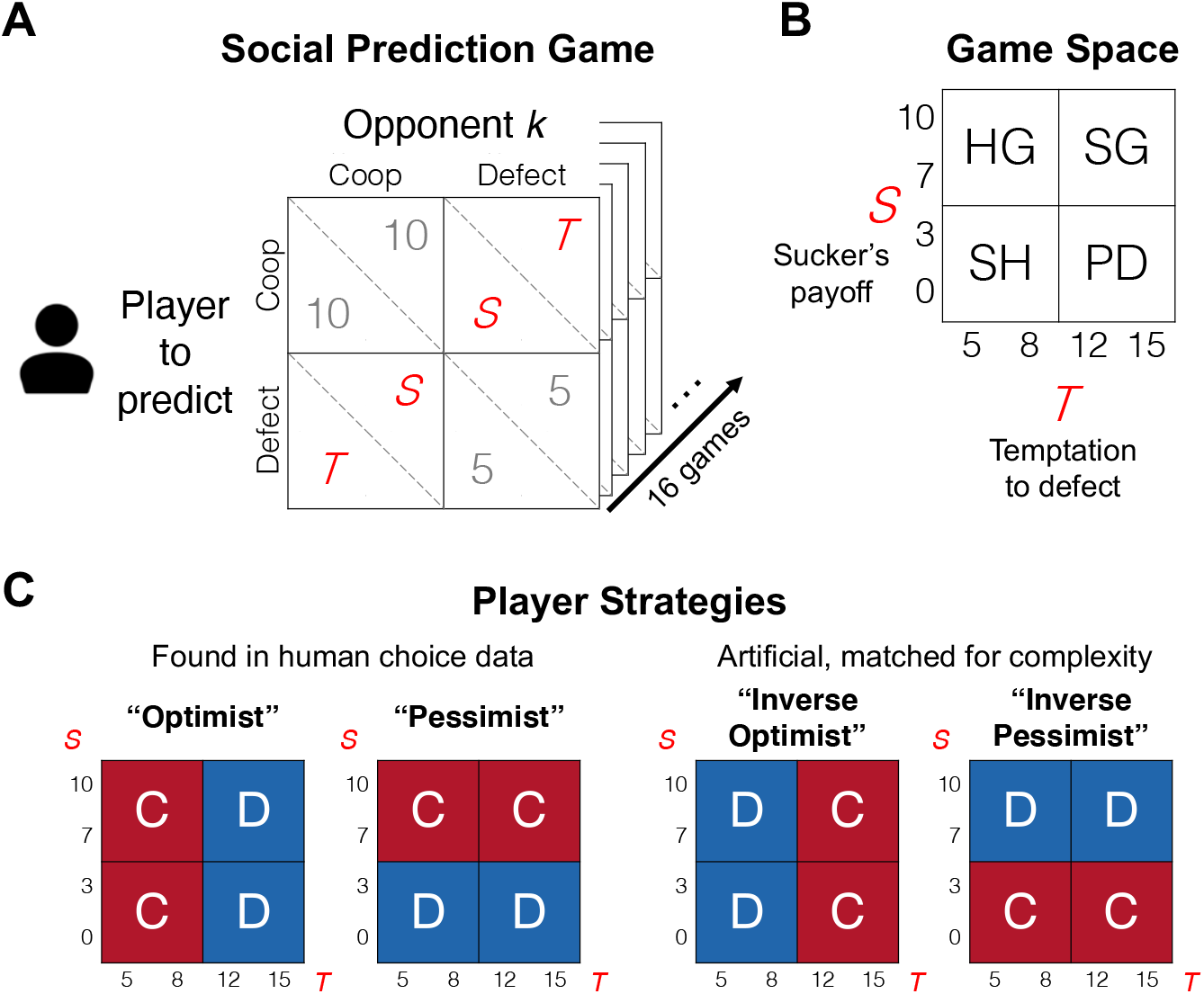
The Social Prediction Game. **A.** On each trial, the social prediction game is presented as a payoff matrix describing the single-shot interaction between a Player (who is tracked across 16 consecutive trials) and anonymous Opponents (who are different on each trial). **B.** The values of S and T vary from trial to trial and are drawn from a 4×4 ‘game space’ such that each trial belongs to one of four canonical economic game types: harmony game (HG), snowdrift game (SG), stag hunt (SH), and prisoner’s dilemma (PD). **C.** Across these different game types, the Player’s true behavior (cooperate, C, or defect, D) depends on the Player’s underlying strategy, which is deterministically programmed in each block of 16 trials.

## Results

In study 1, 150 participants played the Social Prediction Game. The games between Players and Opponents were presented as a 2×2 payoff matrix (Figure 1A) where two of the four possible outcomes yield the same payoffs across all trials: (10,10) for mutual cooperation and (5,5) for mutual defection. The payoffs assigned to the other potential outcomes, labeled S for sucker’s payoff and T for temptation to defect, are drawn from a 4×4 grid (the ‘game space’) such that each of the 16 trials is a new game with a unique (S,T) combination. The varying (S,T) payoff values yield four canonical economic game types which are represented in four quadrants of the (S,T) game space (Fig. 1B). For example, the classic one-shot Prisoner’s Dilemma is found at S < 5 and T > 10, where all players are best-off defecting regardless of their opponent’s choice (i.e. mutual defection is the unique Nash equilibrium)^34,35^. Due to these regularities in game incentives, human choice data reveals several distinct patterns of correlated choice (i.e. strategies) across the 16 games, with each strategy optimizing for a distinct human motive like risk aversion or envy^14^. We programmed the Players to behave in line with one of four deterministic strategies: two found in human choice data at similar rates, namely a greedy strategy labeled Optimist and a risk-averse strategy labeled Pessimist (respectively 20% and 21% of the population^14^), and two not found in human choice data but matched for statistical complexity (i.e., Inverse Optimist and Inverse Pessimist; Fig. 1C).

### Selective generalization of social information

How might participants approach the Social Prediction Game? One possibility is that they expect a Player to simply repeat their past behavior due to stable preferences for cooperation^10^, which amounts to basic reinforcement learning over the response options ‘cooperate’ and ‘defect’ without distinguishing between games. Another possibility is that participants refrain from generalizing across games at all, because each trial is unique. Since all Players cooperate and defect on half the trials, both these strategies would yield on average 50% accuracy in our task. However, observed accuracy was significantly greater (59.1%; two-tailed one-sample *t*-test: *t*(149) = 12.2, *p* < 0.001).

A third possible strategy is naïve statistical learning, whereby participants detect the mapping between S or T and the Player’s choices, e.g. learning that the Inverse Pessimist cooperates when S < 5. This reflects how participants learn latent structure in non-social tasks containing abstract stimuli like colored shapes and fractals^27,28^. If true, task performance should be equal across all Player strategies as each strategy is a step function with a single change point on the S or T dimension (see Figure 1C). However, performance was much higher for human than artificial strategies (Optimist and Pessimist: average accuracy 71.6%; Inverse strategies: 46.5%; two-tailed paired-samples *t*-test: *t*(149) = 22.0, *p* < 0.001; Figure 2A), revealing that naïve statistical learning does not capture learning in this task.

**Figure 2.**
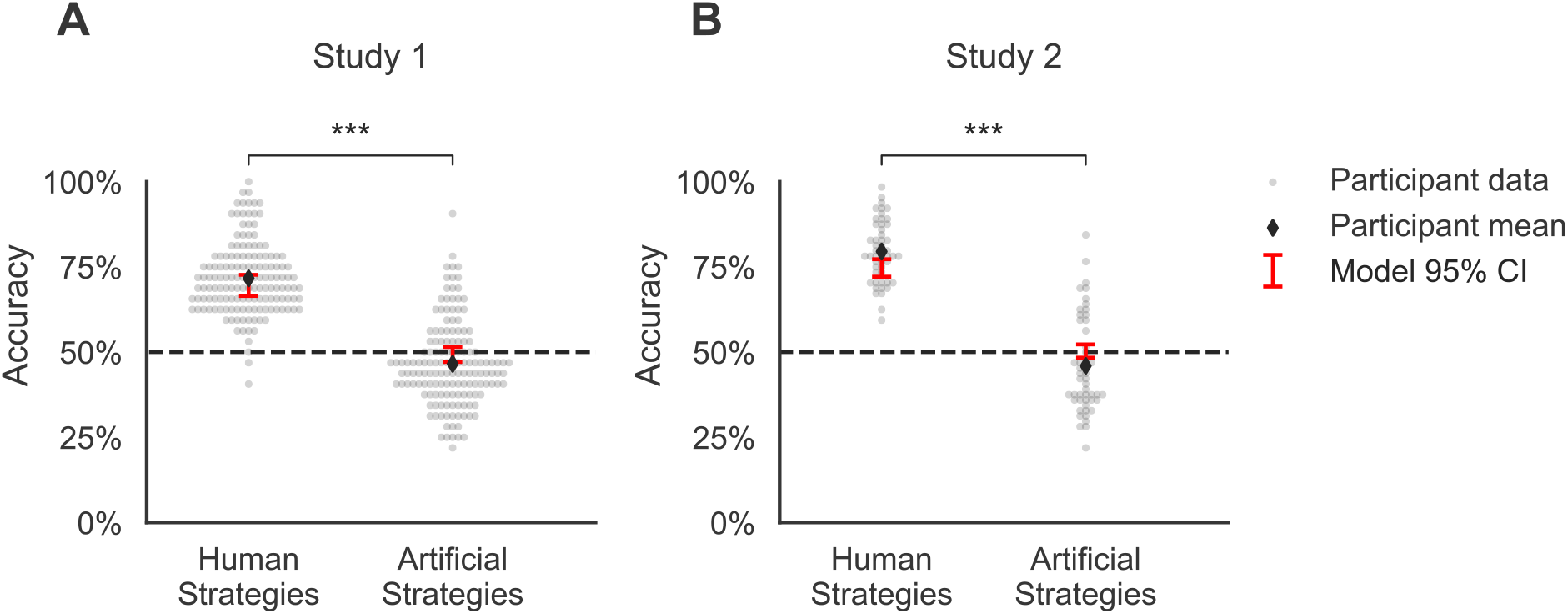
Behavioral results. In studies 1 (**A**) and 2 (**B**), task performance for inferring human strategies (Optimist and Pessimist) was greater than inferring artificial strategies (Inverse Optimist and Inverse Pessimist), and the computational model accurately reproduced this effect. Each gray dot is a participant. Black diamonds indicate average participant accuracy. Red error bars are 95% confidence intervals of average accuracy obtained from simulating the fitted model 1000 times per participant. ***P<0.001.

These findings demonstrate that a) participants selectively generalize information across games by learning a latent structure that links game incentives to choice, and b) participants hold strong prior expectations about the nature of this latent structure, as they were able to predict the choices of the human Players but not artificial ones. We next used computational modeling to illuminate the nature of this latent structure and how participants acquired it when generalizing information about others’ decisions across games.

### Motives guide social structure learning

To determine how participants generalize information across games during social prediction, we built a formal model of structural learning that could capture how participants map the S and T game variables onto choices made. To do this, we leverage recent developments in feature-based reinforcement learning where agents do not learn the value of stimuli per se (e.g. apples or oranges), but rather the decision relevance of stimulus features (e.g. their size or color)^24,29^. We adapted this class of model by allowing it to learn not just over observable task features S and T, but also over unobservable features such as game types or a Player’s motives.

Task behavior was best captured by a model that generalized information across games according to four human motives: Cooperativeness, Greed, Risk aversion, and Nash equilibrium play (*structure by motives*; Figure 3A; see Methods). Model comparison using the Bayesian Information Criterion (BIC)^36^ showed that the average fit of this model was superior to two alternative models: a model that treated each trial as unique (*no structure*; mean ΔBIC = −55.41, Wilcoxon sign-rank test *W* = 7, *p* < 0.001) and a model that generalized according to game type (*structure by game type*; mean ΔBIC = −5.95, *W* = 1641, *p* < 0.001; Figure 3B). Quality checks (Supplemental Information section SI.1) and posterior predictive checks (Figures 2A and 3C) confirmed the robustness of the *structure by motives* model. These results support our hypotheses that participants rely on a feature-based reinforcement learning system that leverages latent motives as a mental scaffold, to generalize what they know about another person across social settings.

**Figure 3.**
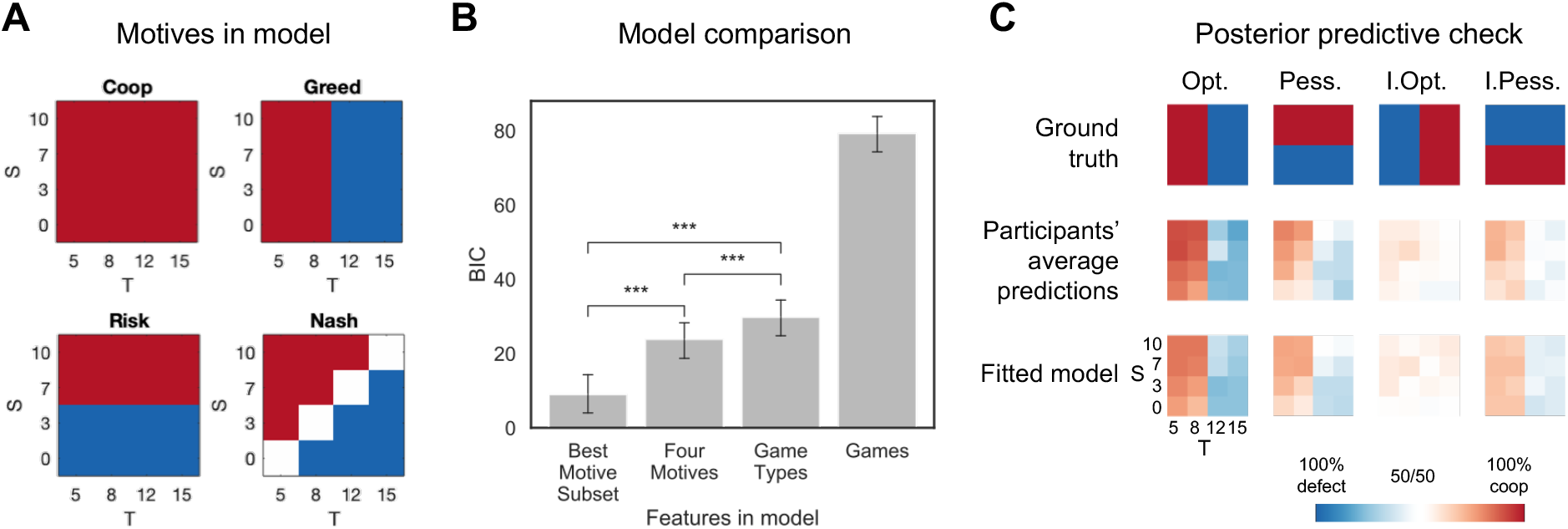
Computational models. **A.** The four motives included in the winning ‘ structure by motives’ model make different behavioral predictions across the S-T game space. **B.** Model comparison for study 1 showed that each participant’s behavior was best described by a model including a subset of the four motives. BIC: Bayesian Information Criterion. Error bars represent bootstrapped 95% confidence intervals. *** p < 0.001 in two-tailed, non-parametric Wilcoxon sign-rank test. **C.** Posterior predictive checks show that the model captures the fine-grained patterns of observed behavior in study 1 as a function of player strategy, S, and T.

### Effort-accuracy trade-off in social structure learning

Our observation of structure learning in a social prediction task poses an interesting puzzle. On the one hand, it is likely that structure learning improves social prediction when facing a variety of latent motives in one’s social environment^12,14^. On the other hand, identifying latent motives is effortful, as it taxes working memory (i.e. updating the predictive value of several motives at once) and requires one to pay attention to many different task features that each inform different motives, such as the values of S and T. To trade off accuracy and effort, participants may therefore consider only a subset of the four motives in their mental models of others’ behavior. To test this, we again fit the motives model to the participants’ prediction data, this time comparing different model versions that each contained a unique subset of motives (i.e. only Cooperativeness, only Greed, Cooperativeness + Greed, and so on). We then ran model comparisons at the subject level, again using BIC as criterion to reward model accuracy and penalize complexity (i.e. the number of motives in a given model).

Confirming our hypothesis, we found that the predictions of all participants were best accounted for by a model that specified only a subset of the four motives rather than by the full model containing all four motives (mean ΔBIC = −14.88, *W* = 0, *p* < 0.001; Figure 3B). 30% of participants considered only a single motive, 50% considered two motives, and 20% considered three motives simultaneously. Participants who built more complex mental models showed superior task performance but also spent more time making their predictions (SI.2), demonstrating the effort-accuracy trade-off of structure learning.

Participants also differed in what motives they considered. 87% of participants considered Greed in their mental model, 51% considered Cooperativeness, 31% considered Nash, and only 21% of participants considered Risk aversion. Accordingly, since Greed drives the Optimism strategy, task performance was much higher during the Optimist block (average accuracy 87.0%) than the Pessimist block (average accuracy 56.3%; two-tailed paired-samples *t*-test: *t*(149) = 14.7, *p* < 0.001). To test what drove these individual differences in motives considered, we compared each participant’s best motive subset to his/her decision strategy (which participants reported during the task instructions). The Cooperativeness motive was more commonly considered in others by participants who had reported their own decision strategy to be cooperative during the task instructions (one-tailed binomial test *p* = 0.010), and the same was true for the Risk aversion motive in participants who had reported a pessimistic strategy (*p* < 0.001). These self-anchoring effects suggest that our own motives shape our constrained mental models of others’ behavior.

### Social structure learning facilitates attentional focus

Although participants with more extensive mental models were slower to make social predictions, this relative time cost decreased between early trials (1-8) and late trials (9-16) (SI.2), reflecting the fact that structure learning involves gradually simplifying complex decision problems^31^. Specifically, it has been suggested that structure learning facilitates an attentional focus on information relevant to the (learned) latent structure of the task^24,26^. We tested this mechanistic hypothesis in study 2, where 50 participants played the same game—behaviorally replicating the findings from study 1 (SI.3)—only this time in the laboratory and while undergoing concurrent eye-tracking. We hypothesized that as participants learned the motives of the Player, they would focus their attention to the task features relevant to those specific motives, namely the temptation to defect T for the Optimist player and the sucker’s payoff S for the Pessimist (Figure 1C).

Confirming our prediction, participants spent a greater percentage of the trial looking at T relative to S in the Optimist block than the Pessimist block (two-tailed paired-samples t-test *t*(49) = 5.11, *p* < 0.001; Figure 4A). To test whether this gaze difference developed as a function of using a structure learning mechanism, we split the Optimist and Pessimist blocks into four mini-blocks of four consecutive trials in which each game type was visited once, and ran a mixed-effects regression with random subject intercepts to test whether Player strategy (Optimist/Pessimist) interacted with mini-block number to predict relative gaze (S versus T). The strategy-mini-block interaction was significant (*F*(1,347) = 16.9, *p* < 0.001) with the greatest shift in attention taking place in the first 4-8 trials (Figure 4B), confirming that the attentional shift between the temptation to defect and the sucker’s payoff was conditional on learning about the Players’ strategies.

**Figure 4.**
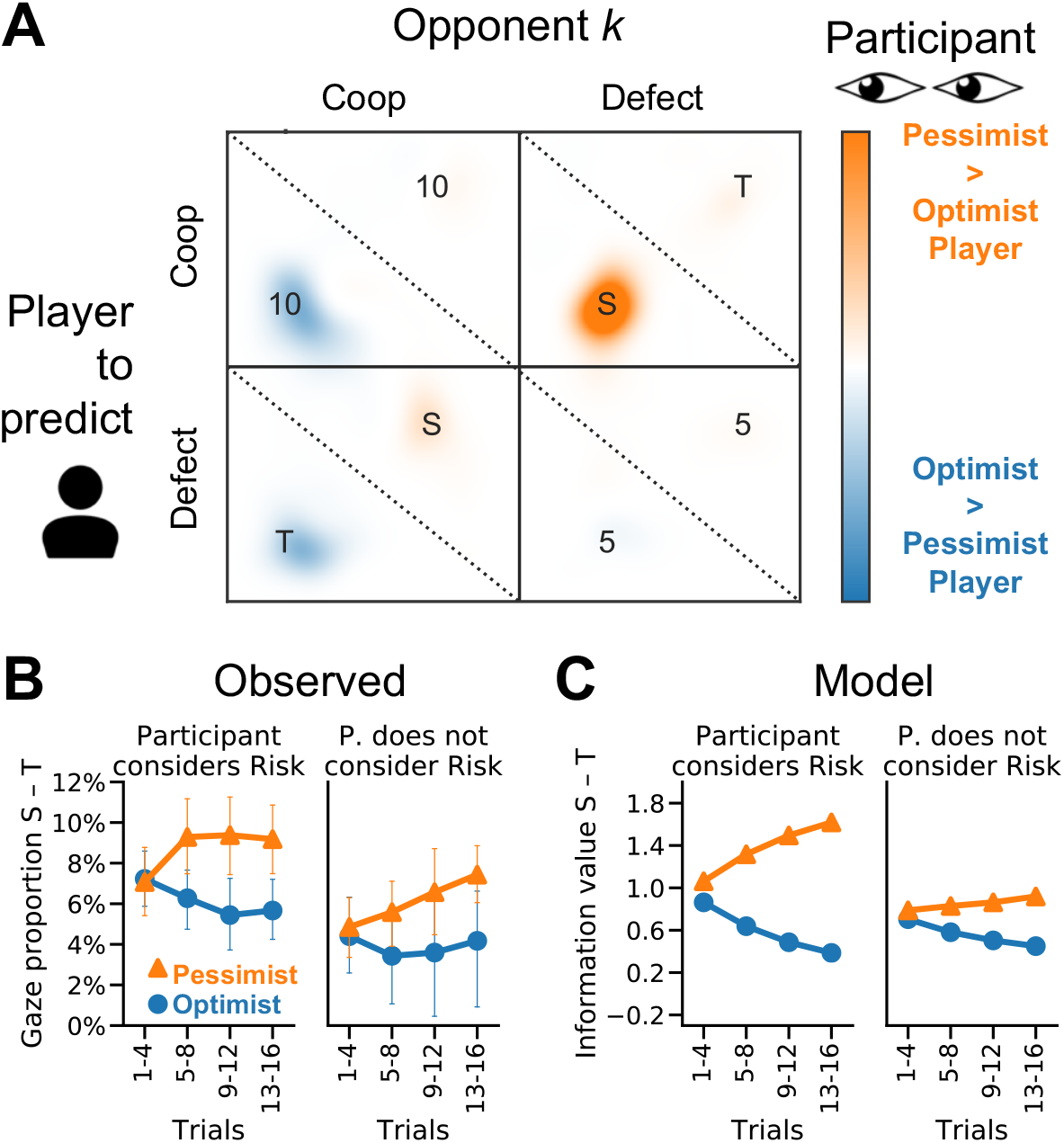
Eye-tracking results for study 2. **A.** Participants selectively looked at information diagnostic of a Player’s strategy, i.e. T for Optimist and S for Pessimist. **B.** Selective attention developed over time as participants learn the Player’s motives. **C.** The computational model was fit only to choice and confidence data, but qualitatively reproduced the participants’ gaze shift over time as displayed in panel B.

The gaze data also validated our computational model, as the model was able to predict gaze over time as a function of the participant’s beliefs about the player’s motives (SI.4; Figure 4C). Finally, participants who shifted their gaze more to the diagnostic information (T for Optimist, S for Pessimist) also performed better at the task (Pessimist block, correlation between relative gaze to S versus T and accuracy: *r*(48) = 0.37, *p* = 0.008; ceiling effect in Optimist block with on average 14 of 16 trials correct), establishing an important role for attention in scaffolding social structure learning.

### Structure learning supports adaptive social choice

In studies 1 and 2, we found that people use a structure learning system to infer the latent motives of others in order to predict which social decisions those others will make. People likely engage in social structure learning—despite the associated effort—because it enables them to strategically adapt their behavior to what others will do. However, deploying strategic social behavior requires a deep and flexible understanding of the underlying latent structure of another’s motives, which can enable a person to generalize another’s motives to contexts that have never been experienced before.

To test this generalization effect to new situations, we carried out a third study in which participants first played a block of the Social Prediction Game where the Player was programmed to be either an Optimist or Pessimist. Afterwards, participants played the Inspection Game^37,38^, playing as the Employer and the Player taking the role of the Employee. In the Inspection Game, the Employee receives a wage from the Employer and chooses to either ‘work’ (which creates revenue for the Employer but is costly for the Employee) or ‘shirk’. Because the Employee receives the wage regardless of whether he or she actually works, shirking has the higher payoff if it goes unnoticed. However, the Employer can choose to pay a cost to inspect the Employee and withhold the wage if the Employee is found shirking. Given these rules, an Optimist Employee—who is motivated by greed—will always shirk to obtain the maximal payoff, while a Pessimist Employee—who is motivated by risk aversion—will always work to avoid being caught shirking (Figure 5A). Conversely, it is best for the Employer to pay for inspecting only if the Employee is likely to shirk. The natural mapping of motives in the Social Prediction Game to the Employee’s behaviors in the Inspection Game create a strong test for whether successful structure learning in one context is adaptively used to behave more strategically in an entirely different context. We predicted that successfully learning the motives of the Player during the Social Prediction Game should lead to paying more to inspect an Optimist than a Pessimist in the Inspection Game.

**Figure 5.**
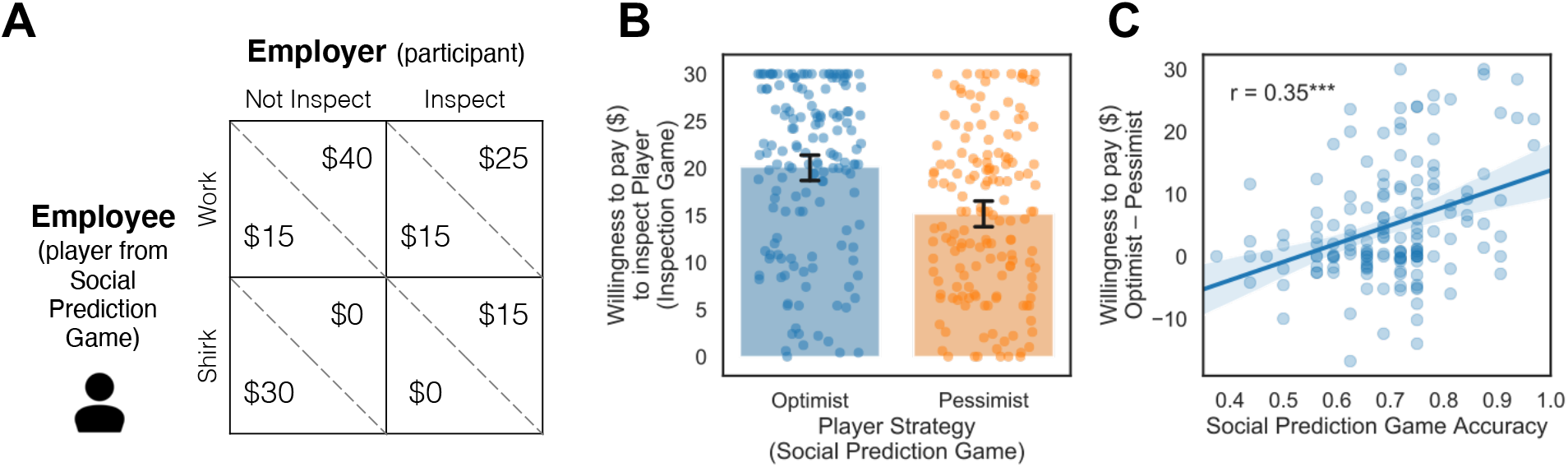
Study 3’s task and results. **A**. Payoff matrix of the Inspection Game, which was played after completing the Social Prediction Game. **B**. Despite an absence of feedback in the Inspection Game, participants paid more to inspect Employees who exhibited Optimistic versus Pessimistic strategies in the preceding Social Prediction Game, revealing generalization of structure across tasks. Each dot is a participant; overlapping dots have darker colors. **C**. The difference in Willingness to pay for inspecting between Optimist and Pessimist is predicted by learning performance on the Social Prediction Game.

Confirming our hypothesis, participants were willing to pay an average of $4.99 more to inspect the Optimist ($20.10) compared to the Pessimist ($15.11; within subject; two-tailed Wilcoxon signed-rank test: W = 2031.5, *p* < 0.001; Figure 5B). This reveals that participants were successful in detecting another’s latent motives in one context and generalizing this information to make more adaptive choices in new, never before experienced situations. Furthermore, a participant’s willingness to pay was mediated by how well they inferred the motives in the Social Prediction Game. The better a participant performed in the Social Prediction Game, the more they paid to inspect Optimists relative to Pessimists (*r*(151) = 0.35, *p* < 0.001; Figure 5C). These costly inspections paid off: The more participants distinguished between Optimistic and Pessimistic Employees in the Inspection Game, the more they earned as an Employer (at $15 default inspection cost: *r*(151) = 0.72, *p* < 0.001).

## Discussion

People routinely predict the behavior of others across a dizzying array of social situations. We have shown that such sophisticated social prediction is achieved through structure learning: participants credit another person’s choices to latent, unobservable motives, such as greed and risk aversion, and use this motive-based abstraction to successfully predict decisions in entirely new interactions with different social tensions. Through structure learning, participants were able to disregard irrelevant contextual cues and attend to information that was diagnostic of the other player’s future actions. Better use of structure learning led to more strategic behaviors in a subsequent competitive decision task with the same player, reflecting the adaptive value of accurate social prediction in everyday life. Together, these findings provide a mechanistic and computational account of social prediction, illuminating how humans adaptively reduce a principal source of uncertainty in our social world^39^: Other people.

This work establishes structure learning^24,25,29,31^ as a critical mechanism for successful social cognition. In our experiments, popular learning models such as reinforcement learning and naïve statistical learning could not account for the selective generalization of social information exhibited by participants. Instead, participants stripped down the complexities of the social exchange, leaving only a few latent dimensions by which to generalize what they had learned to entirely new contexts. This structure learning process was captured by a feature-based reinforcement learning model, originally developed for tracking relevant stimulus features in non-social decision tasks^24–26^, which we adapted to learn over unobservable features of human social engagement such as motives. This establishes feature-based RL as an algorithmic implementation of structure learning in social cognition, in contrast to existing accounts using non-parametric Bayesian models^20,40–42^. In line with the predictions of feature-based RL^24^, response times were slower on early trials, when participants were considering multiple motives as potential causes of the player’s choices, and became faster as participants honed in on a single motive and deployed selective attention to information relevant to that motive. This gradual narrowing of attention allowed participants to limit cognitive resource use while still achieving accurate social prediction. In this way, motive-based structure learning is to social cognition what chunking is to working memory^43,44^: a cognitive adaptation that leverages the inherent structure of the world.

This cognitive adaptation allows us to navigate our complex and evolving social environment by uncovering its latent structure, which is a better basis for generalization than simply attending to context cues or others’ actions. Consider a toy example of a resistance hero who never lies to her parents, but during enemy occupation staunchly denies being part of the opposition to protect her family. A superficial ‘bookkeeping’ of this person’s behavior—she never lies—would fail to predict her actions in the novel context of war, while a deeper understanding of her latent motives—caring for her family—would facilitate accurate behavioral predictions. Similarly, in the Social Prediction Game, Optimist and Pessimist players cooperated and defected equally often (50% of trials), and therefore generalizing only by their observable choices would yield identical (and highly uncertain) predictions in all new games. However, by abstracting and generalizing the Players’ latent motives from the combination of game incentives and choices made, our participants correctly predicted divergent behavior in an entirely new context, the Inspection Game^37,38^. After successfully learning latent structure, participants paid for inspections *only* when it was valuable to do so, thereby earning significantly more money. This intimates that many strategic socio-economic behaviors that rely on predictions about others’ future choices in novel contexts, such as competitive bargaining^45^, market entry^46,47^, and collective action^2,33,48,49^, are scaffolded by a motive-based structure learning mechanism. Given that social motives (e.g., greed) vary between individuals in the population^10,12,14,15,50^, effective inference of these motives in others is likely instrumental in achieving competitive and collaborative goals.

How do people construct and apply abstracted mental models of others’ motives? Our data suggest that attention plays a key role in guiding this process. Attention is a fundamental cognitive mechanism as it affords optimal access to behaviorally relevant information with limited processing capacity^51^. Our findings show how attention supports social prediction. In the Social Prediction Game, as in everyday social interactions, there were multiple cues that could be predictive of another’s behavior, from the player payoffs S and T to the order of the games or even the initials of the player. Structure learning allowed participants to disregard superficial cues and attend to information relevant to the players’ latent motives. Although this process facilitated accurate social prediction with limited effort if the inferred motives were correct, incorrect structure learning caused counterproductive attention on irrelevant information. For example, participants who did not consider risk aversion failed to shift their attention to the sucker’s payoff (S) during the Pessimist block and instead kept looking at the temptation to defect (T), thereby missing out on information predictive of the player’s choices. This suggests that what we can learn about other people is limited by our expectations. Our mental models of others effectively constrain what we pay attention to, causing blind spots that prevent us from properly updating our beliefs and potentially contributing to stereotype^52,53^ and confirmation bias^54^.

Our findings have implications for understanding these social behaviors in the real world. For example, police officers may disproportionally target racial minorities^55^ because of a bias in information search, where inaccurate mental models of minorities’ motives cause police officers to misinterpret behavior or ignore exonerating information. Although future work can directly test this hypothesis, our tasks and models help to disentangle the complementary roles of prior beliefs, information search, and structure learning in social prediction, providing new avenues for understanding failures of social prediction observed in everyday life.

## Methods

### Study 1: Procedure

This experiment (as well as Study 2 and 3) was approved by the Institutional Review Board of Brown University. 150 participants (95 males, 52 females, 3 no response; mean age 35.4 ± 10.0y) participated via Amazon Mechanical Turk (MTurk). All gave written informed consent before starting the experiment. The task was written in Javascript and made accessible using Psiturk 2.3.0^56^. Participants first read instructions and were quizzed to ensure their understanding and filter out potential bots. Participants were then asked to indicate for each game type in the Social Prediction Game how they themselves would choose, from which we estimated the participants’ own decision strategy. They then completed the Social Prediction Game (SPG).

### Task

Participants played 4 blocks of the SPG, each block with a different Player, and were tasked with predicting the choices of this particular Player across 16 consecutive economic games. Players always played single-shot against anonymous Opponents. Each game was presented as a 2×2 payoff matrix (Figure 1A) where the Player and Opponent both have two choices: cooperation and defection. In the task, these choices were labeled by arbitrary color words (blue, green, etc.) whose mapping to cooperate and defect changed on every task block.

The games varied on two features central to social interactions: Risk of cooperating (here operationalized as the sucker’s payoff, S) and temptation to defect (T) (Figure 1B). At T < 10 and S > 5, the games fall under a class of Harmony Game, where each player’s payoff-maximizing action aligns with the jointly payoff-maximizing action and thus no conflict arises except through potential envy^57^. At T > 10 and S > 5 the games are classified as Snowdrift Games (also known as Volunteer’s Dilemmas), which are anti-coordination games where unilateral defection is preferable to mutual cooperation, but mutual defection yields the smallest payoff for all^58^. At T > 10 and S < 5 lie the Prisoner’s Dilemma games, which are characterized by a high temptation to defect even if one’s opponent defects as well, and cooperation is risky as unilateral cooperation yields the lowest possible payoff^34^. At T < 10 and S < 5 the games are Stag Hunts, in which mutual cooperation yields the highest payoff for both, but cooperation is risky as unilateral cooperation is met with the lowest payoff^59^.

The task of the participant was to indicate, on each trial, what they believed the Player would choose to do in the current game, and to rate their confidence in this prediction on an 11-point scale from 0% to 100% (10% increments). They received feedback on every trial indicating whether their prediction was correct or not, and earned a $0.01 bonus for every correct trial. At the end of 16 trials (1 block), the participants self-reported what they believed the Player’s strategy was using a free-response answer box. After four blocks, the total earned bonus was presented to the participant and added the base payment. The participant was then taken to a survey hosted on Qualtrics to finish the experiment.

### Computational models

We built three versions of the model with different types of features that each reflect a different psychological model about how participants might carve up the 4×4 game space in order to generalize information across trials.

Model 1 (*structure by motives*) learns over four features representing psychological motives that span the entire game space. This reflects the hypothesis that participants realize that the Player’s behavior across all games is driven by overarching motives, which is used to generalize learning across games to improve predictions. Model 1 can support sophisticated generalization across the game space whereby cooperation in one game type predicts defection in another—a common pattern in human choices caused by the fact that consistently adhering to one motive can lead to different choices in different games^14,33^. As features in this model, we included four motives whose influence on behavior in economic games is well-documented: Cooperativeness (which always predicts Cooperate)^60^, Greed (which maximizes the maximal payoff)^5^, Risk aversion (which maximizes the minimal payoff)^61^, and Nash (which plays the Nash equilibrium)^35^ (Figure 3A). Note that the model does not assume that these motives drive the participant’s own choices, but rather that the participant might consider these motives as drivers of the other Players’ decisions.

Model 2 (*no structure*) learns over 17 features: One for each unique (S,T) combination and one ‘intercept’ feature that spans the entire game space. It can thus generalize information across all games equally and treat each game as unique, but it cannot selectively generalize within a subset of games. It reflects the hypothesis that participants generalize learning about a Player only in the coarsest way possible—across all games equally—without realizing how several different games might elicit the same, or different, decisions from a Player.

Model 3 (*structure by game types*) learns over five features: One for each game type (i.e. quadrant in the game space) and one intercept. This model reflects the hypothesis that participants recognize that behavior is consistent within a game type (e.g. stag hunt) and thus benefits from grouping learning experiences by game type. Since Player behavior is fully consistent within game type, this strategy could be very successful in the current task.

In each of our three candidate models, predictions for the Player’s choice are made based on the weighted average predictions of each included model feature (i.e. games, game types, and motives; see Figure 3A-C):

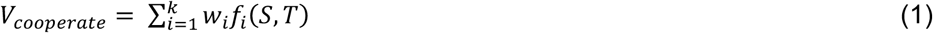

Where *f*_*i*_(*S*, *T*) is the feature’s prediction at the current S and T values, expressed as 1 for Cooperate and −1 for Defect; *w*_*i*_ is the associated feature weight; *i* is the feature number; *k* is the total number of features, and *V*_*cooperate*_ is the ‘value of predicting cooperate’. We then apply a softmax decision rule to produce the model prediction:

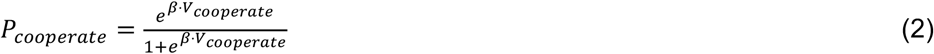

Where *β* is the inverse temperature parameter of the softmax function and *P*_*cooperate*_ represents the likelihood that the Player will cooperate given the current state of the model. During learning, the Player’s actual choice is compared to the model’s prediction to compute a prediction error *PE*:

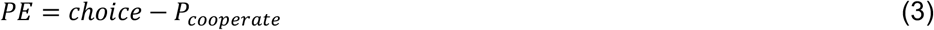

Where *choice* is 1 if the Player cooperated and 0 if the Player defected. Therefore, the prediction error is >=0 if the Player Cooperated and <= 0 if the Player Defected. This sign difference is used in the next step, where each feature weight *w*_*i*_ is updated based on the prediction error, the learning rate *α*, and the direction of the prediction, for all *i* from 1 to *k*:

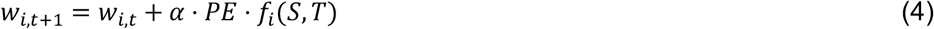

Where *f*_*i*_(*S*, *T*), as before, is 1 for Cooperate and −1 for Defect. Combining this sign with the sign of the prediction error allows the model to increase a feature weight if the prediction of the associated feature was correct (i.e. *f*_*i*_ = 1 and *PE* > 0 or *f*_*i*_ = −1 and *PE* < 0), and conversely to decrease the weight if the feature was incorrect (when *f*_*i*_ and *PE* have opposite signs).

### Model fitting

The feature weights are reset to their *a priori* values at the start of each task block (i.e. predicting each new Player). These prior weights are free parameters that are determined when fitting the model to each participant’s data, along with the learning rate *α* and the softmax inverse temperature *β*. Model 1 thus has six free parameters (four feature weights, *α*, and *β*), Model 2 has 19, and Model 3 has seven.

To fit the free parameters, we defined the objective function as the sum of squared errors between the model prediction and the participant’s prediction weighted by confidence. Confidence-weighted predictions *x* are computed as follows:

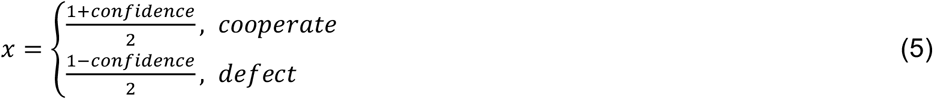

For example, if the participant rated 70% confidence that the Player would defect, the confidence-weighted prediction is (1 – 0.7) / 2 = 0.15. To compute the sum of squared error of the model, we compared the confidence-weighted prediction to the model prediction:

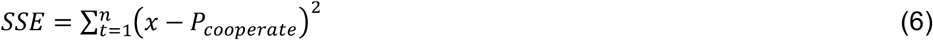

Where *t* indicates trial, *n* indicates the total number of trials (64 in study 1, 128 in study 2), and *SSE* indicates the sum of squared model error. We used the Matlab function *fmincon* to find the parameter combination that minimizes *SSE*. Finally, we computed BIC to penalize model fit by model complexity (i.e. number of free parameters), using the assumption that model error was normally distributed:

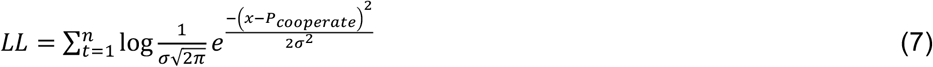

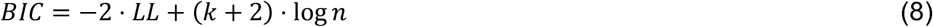

Where *σ* is the standard deviation of the model errors *x* − *P*_*cooperate*_, *k* is the number of feature weights in the model (*k* + 2 thus represents the number of free parameters), and *n* is the number of observations (trials). BIC is an appropriate choice of model selection metric, as it allowed for reliable model selection in synthetic data (SI.1).

### Study 2: Procedure

50 participants (38 women, 12 men; mean age 22.2 ± 7.3y) played 8 blocks of the Social Prediction Game in the laboratory while undergoing concurrent eye-tracking. All participants gave written informed consent before the start of the experiment. Each participant played two blocks with each Player type (Optimist, Pessimist, etc.) in pseudorandom order such that each Player type occurred once in the first four blocks and once in the second four blocks. Each Player was labeled with a unique set of initials, regardless of strategy. Participants took a break between block 4 and 5. The task was generated using the Psychophysics Toolbox for Matlab. Otherwise, the task and computational model used in study 2 were identical to those in study 1.

### Eye-tracking

Eye tracking data was collected using a SensoMotoric Instruments (SMI; Teltow, Germany) iView X RED. Participants were seated comfortably and upright in a straight-backed chair at 50-70 cm from the screen, and were instructed to move as little as possible. The Social Prediction Game was preceded by a proprietary SMI eye tracker calibration and validation procedure in Matlab, which was repeated if necessary until the deviation in visual angle between the validation target and the recorded gaze fixation was below 2 degrees. Average deviation after calibration (X,Y) was (0.65°, 0.86°) for the left eye and (0.74°, 0.88°) for the right eye. The calibration and validation procedures were repeated after the break before task block 5.

Gaze data were preprocessed by converting the raw data to fixations by grouping data points based on spatial and temporal distance using SMI software with default settings. Next, all fixations outside of the bounding box of the rectangular payoff matrix on the screen were excluded. Circular regions of interest (ROIs) were then defined with 100 pixel radius around the center of each of the eight numbers in the payoff matrix. We then computed the relative gaze to each ROI by dividing the summed fixation duration in that ROI by the total fixation duration in the entire payoff matrix. We computed relative gaze to each ROI per eye and then averaged over the two eyes to obtain the overall relative gaze to each ROI. For the S-T difference scores, we subtracted relative gaze to T (summed over both T ROIs on the screen) from relative gaze to S (summed over both S ROIs).

### Study 3: Procedure

153 participants (92 males, 61 females; mean age 37.6±10.4y) gave written informed consent before starting the experiment. They first played one block of the Social Prediction Game and then a block of the Inspection Game (IG), with the same Player. Participants then played another block of the Social Prediction Game and Inspection Game with a different Player. For both Players, they could thus use what they had learned about that specific Player in the Social Prediction Game to improve their decision-making in the Inspection Game. Critically, the Inspection Game is structurally distinct from all games included in studies 1 and 2, as the game is not symmetric (e.g. mutual cooperation yields unequal payoffs). Moreover, whereas our participants only observed other Players in the Social Prediction Game, they actively participated in the Inspection Game and their monetary reward was yoked to this game’s outcomes. The Inspection Game is thus a strong test for selective generalization of learned information about the Player to a novel task.

The two Players were programmed as either Optimist or Pessimist in random order. These two strategies were identical to study 1 and 2 for the SPG; for the Inspection Game the Optimist always shirks and the Pessimist always works. Since there was no feedback during the IG, decision-making was solely driven by what the participant had learned during the SPG. On each round of the IG, the participant could earn money from 1) a base pay of $30 and 2) the revenue generated by a working Employee (but not a shirking one) diminished by 3) the wage paid to the Employee and 4) the cost of an inspection (only if chosen). Therefore, the Inspection Game provides an incentive for the Employer to Inspect in order to avoid paying a wage unnecessarily, but only if the Employee is shirking. The participant’s earnings of one randomly selected round of the Inspection Game were paid out as a bonus with $1 in the game converted to $0.01 for the participant.

At the start of each IG block, the participant self-reported how likely they believed it was that the current Player (indicated by initials) would be working in the Inspection Game. Participants rated the Optimist player at 41.3% likely to work and Pessimist at 54.8%, demonstrating that inferences about others’ motives were explicitly available. During the IG, a staircase procedure determined the participant’s willingness to pay for Inspection on the current round by raising the cost of inspecting if the participant had chosen to inspect on the previous trial, and lowering it if the participant had chosen not to inspect. The willingness to pay (i.e. the indifference point) was computed by averaging the cost of inspecting across the last 5 of the 15 trials of the staircase procedure. Additionally, at the end of each IG block, the participant self-reported how much they would be willing to pay for inspection (between $0 and $30).

### Software and code availability

Low-level behavioral data analysis (e.g. computing averages, running t-tests and one-way F-tests) was carried out in Python 3.7.4 using the packages *Numpy* 1.17.2, *Pandas* 0.25.1, and *Scipy* 1.3.1. Figures were created using *Matplotlib* 3.1.1 and *Seaborn* 0.9.0 for Python. Mixed-effects regressions were carried out in *R* using the packages *lme4* 1.1-21 and *lmerTest* 3.1-1. Computational modeling was performed in Matlab R2019b using the *fmincon* function from the Optimization Toolbox and custom code. All code will be made available online upon publication.

## Acknowledgements

We thank Anxo Sánchez for sharing experimental data from ref. ^14^.

## Supplemental Information

### Supplemental Results

#### SI.1. Model quality checks

First, we confirmed that the model fit of Model 1 (*structure by motives*) was superior to the best alternative (*structure by game types*) for the majority of participants (113 out of 149; excluding one outlier who had extremely poor model fit due to always reporting 100% confidence), indicating that the mean BIC difference reported in the main text was not driven by a small number of outlier subjects but rather reflected a consistent effect throughout the dataset. The reduced motives model (*best subset of motives*) was superior to *structure by game types* for 144 out of 149 participants and superior to *four motives* for all 149 participants.

Second, we checked whether our set of four canonical motives was appropriate for describing the motives inferred by our participants. To this end, we generated 5000 sets of pseudo-motives by shuffling the four canonical motive features as defined across the 16 S-T ‘tiles’ in the S-T game space. That is, for each pseudo-motive set we included 1) a feature that predicts cooperate in each game, i.e. identical to the Cooperativeness motive; 2) two features that predict cooperate in 8 contiguous S-T coordinates in the S-T game space, thus retaining the spatial properties of the Greed and Risk aversion motives; 3) a feature that predicts cooperate in 6 contiguous S-T coordinates, 50/50 cooperate/defect in 4 S-T coordinates adjacent to the cooperate coordinates, and defect in the remaining 6 coordinates. Some examples of pseudo-motive sets are given in Supplemental Figure 1A. Results showed that our 4 canonical motives yielded a superior model fit (mean SSE across participants = 4.10) to the 5000 pseudo-motive sets (Supplemental Figure 1B), confirming that we likely did not overlook important motives or alternate bases for generalization when evaluating our winning model.

**Figure S1.**
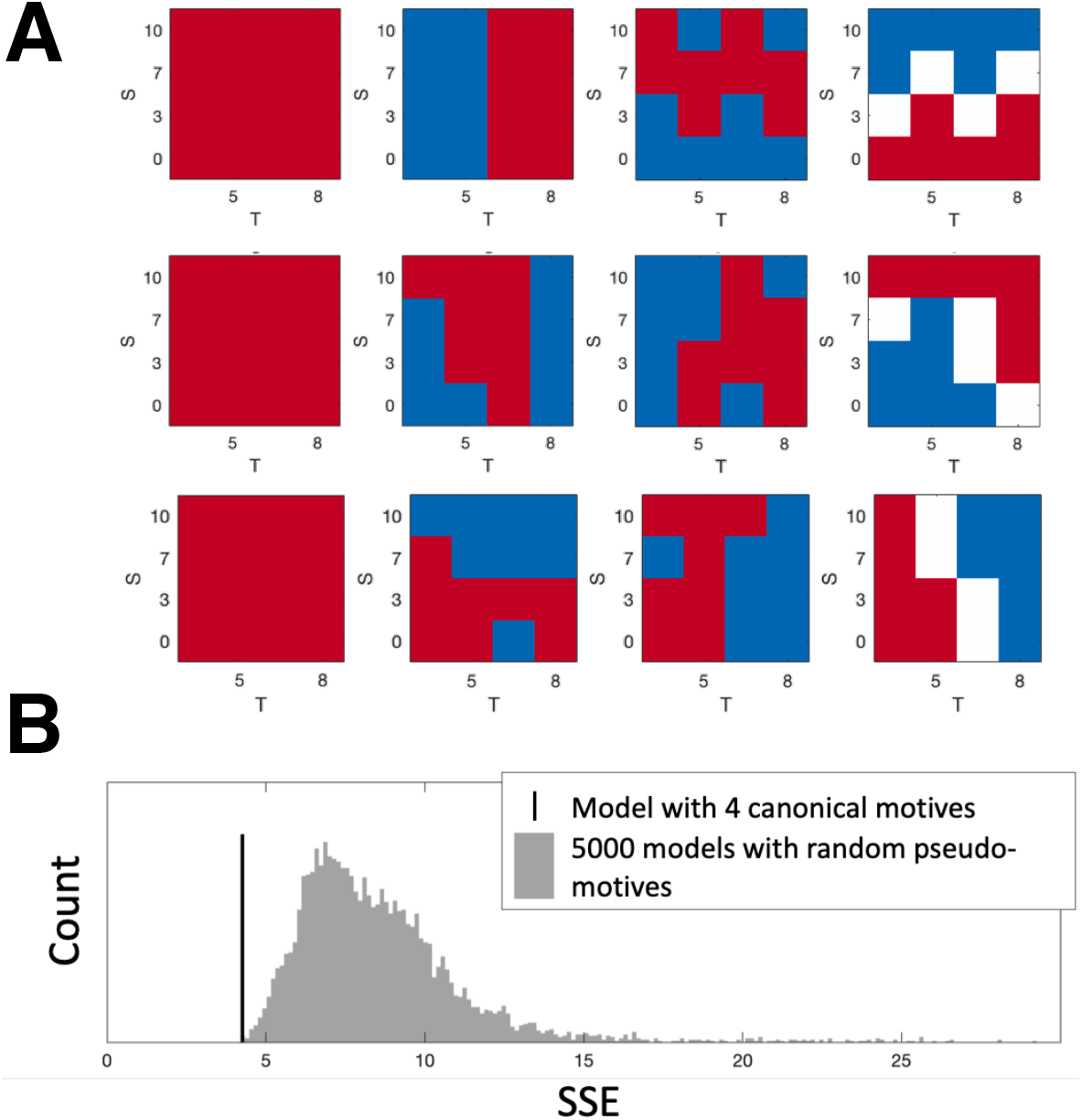
Comparison between Model 3 with 4 canonical motives (Cooperativeness, Greed, Risk aversion, Nash) and 5000 randomly generated pseudo-motives (which are the 4 canonical motives shuffled in the S-T game space). **A**. Three examples of randomly generated pseudo-motives. **B**. Histogram of model error when populating the computational model with the pseudo-motives (grey bars) versus the four canonical motives described in the main text (black vertical line). SSE: sum of squared error.

Third, since our analyses rest on a model selection procedure across subjects as well as within subjects, we tested whether our models could in fact be distinguished from one another with the current experimental design. To this end, we ran a model recovery procedure^62^ where we generated 300 pseudo-datasets by simulating each model 100 times, following the experiments (i.e. trial order) of the first 100 participants in our study. We then fit each of the three candidate models to each pseudo-dataset, counting the number of times that the winning model mirrored the data-generating model. Results are displayed in the confusion matrix in Supplemental Figure 2. Inverting the recovery results shows that if the best-fitting model for a given dataset was ‘structure by motives’ (Model 3 in the main text), there was a 73% chance that this was in fact the true data-generating model, while there was a 20% chance that the true model was Model 2 (structure by game types). The reverse error (i.e., the ‘game types’ model winning while ‘motives’ actually generated the data) was less likely at 6%. Therefore, even when accounting for this potential confusion between models, Model 3 outperformed Model 2 for the majority of participants. The model recovery results thus show that the models were indeed distinguishable in the current experimental design, and that the ‘structure by motives’ model can be reliably interpreted as the winning model.

**Figure S2.**
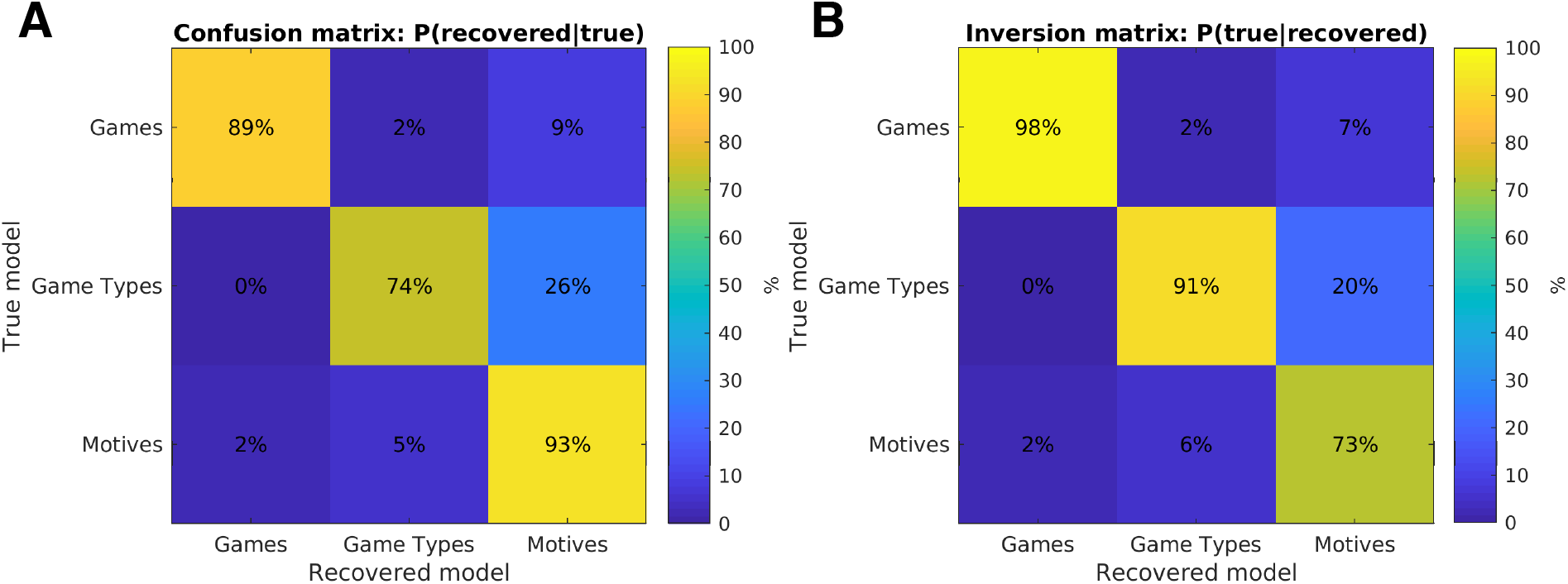
Model recovery results. **A.** Model confusion matrix shows the probability that the data-generated model also wins in model selection using the Bayesian Information Criterion. **B.** Model inversion matrix shows the probability that the recovered model is the true data-generating model. Together, these matrices indicate that the ‘structure by motives’ model is distinguishable from the alternative models given the current experimental design.

#### SI.2. Effort-accuracy trade-off

There were significant differences in task performance between the groups of participants with different numbers of motives in their mental model of the Players (*F*(2,147) = 10.5, *p* < 0.001), with participants who considered three motives exhibiting better performance than those who considered two (two-sided two-sample t-test *t*(119) = 2.24, *p* = 0.027), and two motives yielding better performance than one (t(104) = 2.93, *p* = 0.004; Supplemental Figure 3A).

Although considering more motives was associated with better performance in the SPG, building and updating a complex model of others’ choices comes at a cost. Our model predicts that in early stages of a task block, when the participant does not yet know the strategy of the Player, a complex model (i.e. many motives considered) requires one to evaluate multiple predictions at once. In line with this prediction, average log response times were significantly longer on trials 1-8 than on trials 9-16 (Wilcoxon sign-rank test *W* = 1245, *p* < 0.001).

Moreover, some motives require more information than others. For example, if the participant believes that Players are solely driven by the tendency to cooperate, one does not need to inspect the payoff matrix of the current trial to make the ‘cooperate’ prediction. On the other hand, to predict the actions of a Player who plays the Nash equilibrium, one needs to gather information about both the sucker’s payoff (S) and the temptation to defect (T). To predict the Greed and Risk aversion motives, one only needs one parameter (T or S, respectively). Since information sampling takes time, we predicted that participants who needed to sample S and T would take longer than participants who did not. However, this effect should be specific to early trials of a task block, where participants have not yet narrowed down their mental model of the Player. To test this, we reasoned that participants who considered the Greed motive would focus on T, participants who considered the Risk motive would focus on S, and participants who considered both Greed and Risk and/or the Nash motive would need to attend to both S and T. We thus compared log response times on trials 1-8 of each block for participants who needed to sample *S and T*, *S or T*, or *neither S nor T*. Response times were significantly different (*F*(2,147) = 4.64, *p* = 0.011) for each of these three groups of participants, with attention on *S or T* being associated with slower RTs than not attending to either *S nor T* (Mann-Whitney *U* = 2, *p* = 0.010) and a trend for *S and T* being slower than *S or T* (*U* = 2309.5, *p* = 0.077; Supplemental Figure 3B). As predicted, the response time difference between *S and T* and *S or T* was significantly smaller at the end of a block than at the beginning (two-tailed two-sample t-test on early-late trial difference scores *t*(147) = 2.10, *p* = 0.037). This suggests that as participants learn about the motives of the Player, they limit their attention to task features that are relevant to those specific motives—a hallmark of structure learning^24,25,31^.

**Figure S3.**
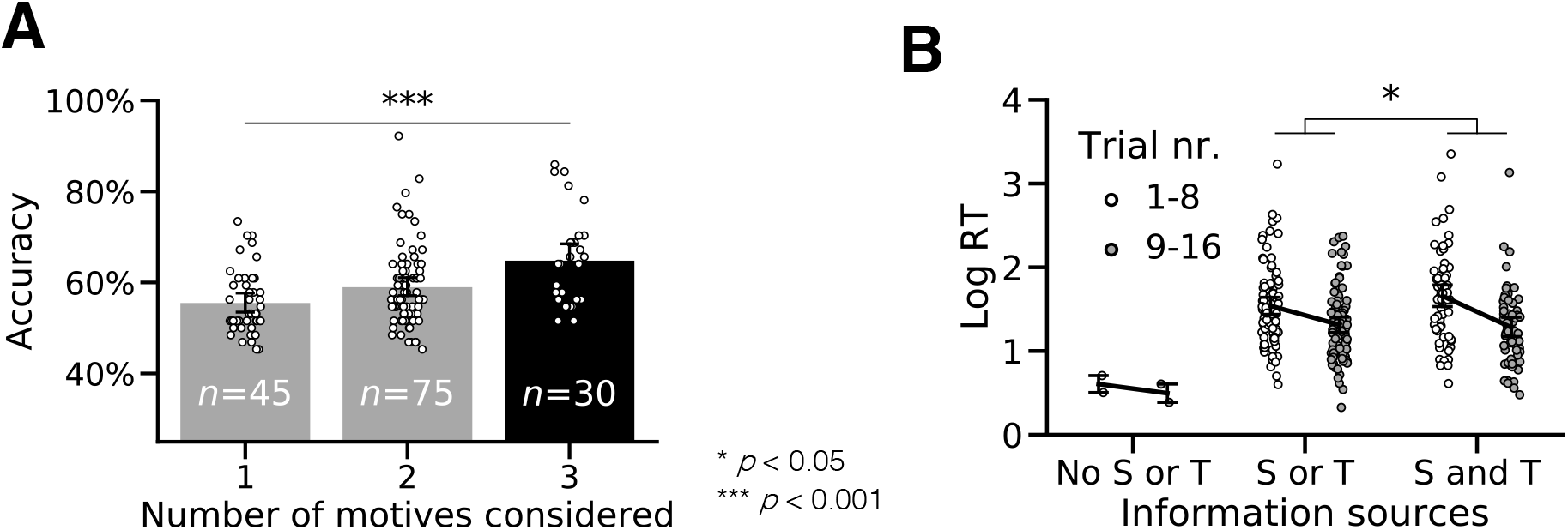
**A.** Accuracy in study 1 as a function of the number of motives considered (i.e. the number of motives included in the participant’s best-fitting model). **B.** Response times in study 1 as a function of the amount of information sampled by participants, based on the motives included in their best-fitting model.

#### SI.3. Replicating study 1’s behavioral effects in study 2

In study 2, as in study 1, participants performed better than chance at the Social Prediction Game (average accuracy 62.7%, two-tailed one-sample t-test against 50%: *t*(49) = 47.5, *p* < 0.001), and were better able to predict human compared to artificial strategies (two-tailed paired-samples t-test *t*(49) = 16.9, *p* < 0.001; Figure 2B). The *structure by motives* model again outperformed the alternative models (mean ΔBIC *motives* – *no structure* = −90.7, *W* = 0, *p* < 0.001; mean ΔBIC *motives* – *game types* = −15.6, *W* = 13, *p* < 0.001) with a subset of the four motives fitting data better than the full model for 49 out of 50 participants (mean ΔBIC = −11.7, *W* = 0, *p* < 0.001).

#### SI.4. Mental models predict gaze

Having confirmed key theoretical predictions about visual attention in social structure learning, we used eye-tracking to test the computational model developed in Study 1. If accurate, this model should be able to predict not just our participants’ behavioral predictions and confidence ratings, but also individual differences in gaze. One way to test this is to compare gaze between participants who did, or did, not consider the Risk motive as driving the Players’ choices, as this motive was considered by about half (58%) of the participants (Cooperativeness: 26%; Greed: 82%, Nash: 56%). We hypothesized that participants whose best-fitting model did not incorporate Risk would focus less on the sucker’s payoff of the games. Confirming this, participants who considered Risk looked significantly more at S (the risk aversion motive) relative to T than those who did not (two-tailed two-sample t-test *t*(49) = 2.04, *p* = 0.047; Figure 4B). This group effect was mirrored with our computational model in a posterior predictive check (Figure 4C).

To provide a strong test of our computational model, we hypothesized that we could predict relative gaze to S and T at the trial level from the computational model fitted only to predictions and confidence ratings. To this end, we computed how informative it would be for a participant at a given trial to look at S or T given the participants’ current beliefs about the Player’s motives. On each trial, we first computed the model prediction *P_cooperate_* for each possible combination of S and T (i.e. the conditional distribution), and compared this to the marginal distribution over S or T using Kullback-Leibler (KL) divergence. For example, consider the case where the model holds that a participant currently believes that the Player is strongly motivated by Greed, a strategy whose choices are determined by the value of T (temptation to defect). In this case, the marginal distribution over T will be very similar to the full conditional distribution over S and T, since the model prediction is dominated by T. This means that the model prediction’s probability distribution will not change much if the participant looks at S, while looking at T will have a big impact on the probability distribution. This change (KL divergence) in probability distribution can thus be interpreted as the value of information associated with S and T, computed as follows:

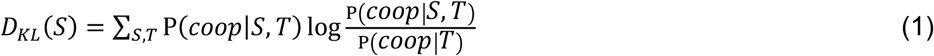

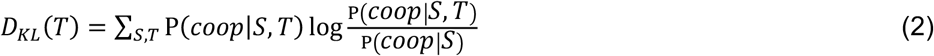

Where *D_KL_(S)* indicates KL divergence of the change in probability distribution upon looking at S, *P(coop|S,T)* denotes the conditional distribution, and *P(coop|S)* denotes the marginal distribution over S. We then predicted relative gaze to S versus T per trial by computing the information value difference *VOI_T-S_* between the two variables:

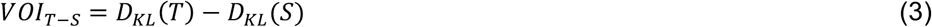

Here, a *VOI_T-S_* of 1 means that given the current belief state of the model, it is only valuable to look at T (e.g. if the model is highly certain that the Player is an Optimist) while inspecting S is only valuable when *VOI_T-S_* is −1.

Using this metric of information value, we were able to predict trial-by-trial gaze of the participants by correlating trial-wise *VOI_T-S_* as generated by the model to the observed relative gaze difference between T and S. The correlation was significantly positive across the 50 participants (two-tailed Wilcoxon sign-rank test of correlation coefficients: *W* = 186, *p* < 0.001; Supplemental Figure 4A). An example participant is shown in Supplemental Figure 4B. This result could never be achieved by a model learning over games (Model 2) or game types (Model 3), as these alternative models assign equal information value to S and T on every trial and therefore cannot account for gaze shifting over time between S and T. These results constitute an out-of-sample test of our computational model, as the model was trained on behavioral data but tested on gaze, and thus provide strong confirmatory evidence that the motives we infer in others drive visual attention in social structure learning.

**Figure S4.**
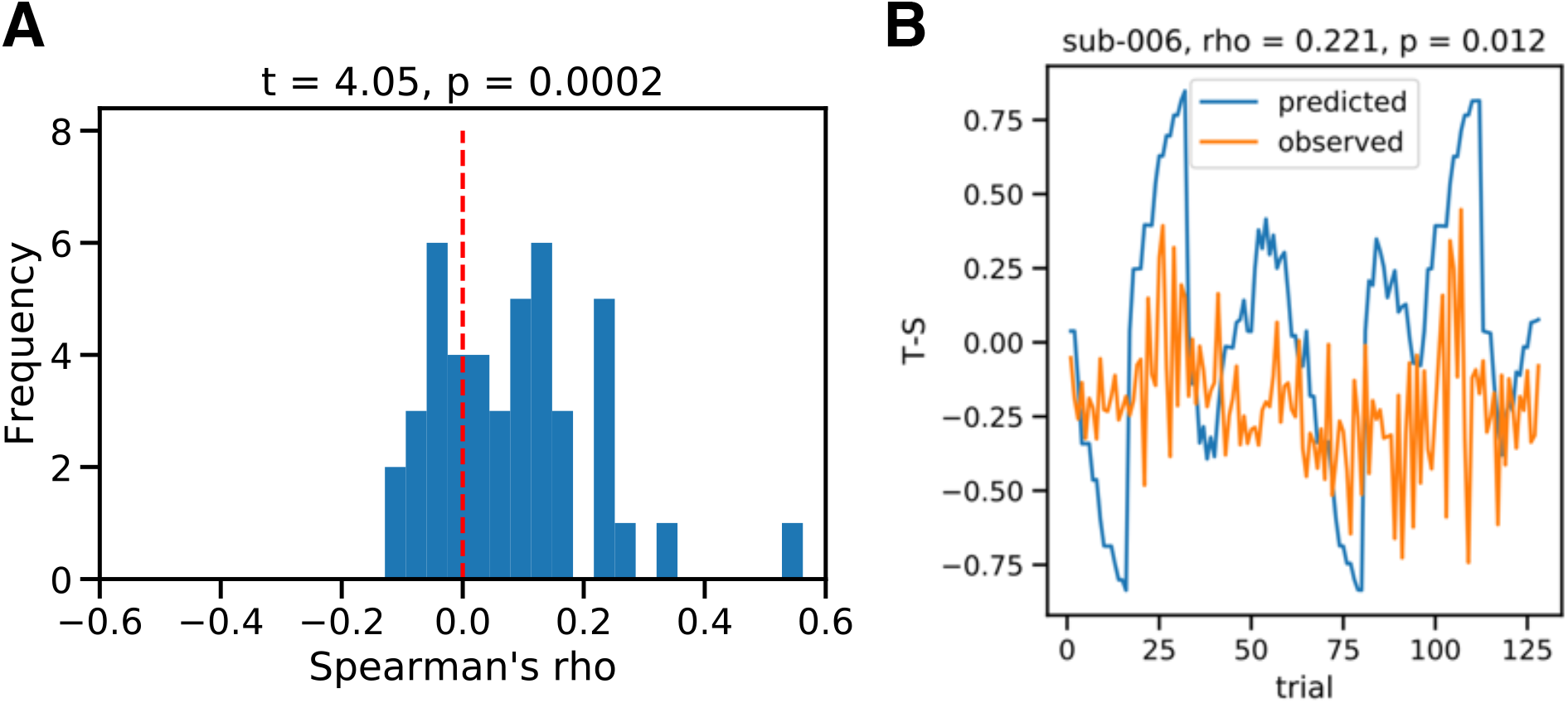
Predicting gaze from the model. **A.** The Spearman correlation between trial-by-trial gaze predicted from the model and observed gaze was significantly positive across the sample. **B.** Data from one example subject illustrate how the model-predicted gaze (blue line) tracks the observed gaze (orange line) trial-by-trial.

## References

1. Stewart, A. J. et al. Information gerrymandering and undemocratic decisions. Nature 573, (2019).

2. Hauser, O. P., Rand, D. G., Peysakhovich, A. & Nowak, M. A. Cooperating with the future. Nature 511, 220–223 (2014).

3. Fraser, C., Riley, S., Anderson, R. M. & Ferguson, N. M. Factors that make an infectious disease outbreak controllable. Proc. Natl. Acad. Sci. U. S. A. 101, 6146–6151 (2004).

4. Tenenbaum, J. B., Kemp, C., Griffiths, T. L. & Goodman, N. D. How to grow a mind: Statistics, structure, and abstraction. Science (80-.). 331, 1279–1285 (2011).

5. Seuntjens, T. G., Zeelenberg, M., Van De Ven, N. & Breugelmans, S. M. Dispositional greed. J. Pers. Soc. Psychol. 108, 917–933 (2015).

6. Fehr, E. & Schmidt, K. M. A theory of fairness, competition, and cooperation. Q. J. Econ. 114, 817–868 (1999).

7. Bolton, G. & Ockenfels, A. ERC: A theory of equity, reciprocity, and competition. Am. Econ. Rev. 90, 166–193 (2000).

8. Weber, E. U., Blais, A.-R. & Betz, N. E. A Domain-specific Risk-attitude Scale: Measuring Risk Perceptions and Risk Behaviors. J. Behav. Decis. Mak. 15, 263–290 (2002).

9. Kahneman, D. & Tversky, A. Prospect Theory : An Analysis of Decision under Risk. Econometrica 47, 263–292 (1979).

10. Peysakhovich, A., Nowak, M. A. & Rand, D. G. Humans display a ‘cooperative phenotype’ that is domain general and temporally stable. Nat. Commun. 5, 1–8 (2014).

11. Van Lange, P. A. M. The pursuit of joint outcomes and equality in outcomes: An integrative model of social value orientation. J. Pers. Soc. Psychol. 77, 337–349 (1999).

12. van Baar, J. M., Chang, L. J. & Sanfey, A. G. The computational and neural substrates of moral strategies in social decision-making. Nat. Commun. 10, 1483 (2019).

13. Driessen, J., Van Baar, J., Sanfey, A., Glennon, J. & Brazil, I. Moral strategies and psychopathic traits. Revis.

14. Poncela-Casasnovas, J. et al. Humans display a reduced set of consistent behavioral phenotypes in dyadic games. Sci. Adv. 2, e1600451 (2016).

15. Van Baar, J. M., Klaassen, F. H., Ricci, F., Chang, L. J. & Sanfey, A. G. Stationary distribution of moral strategies in a population. PsyArXiv (2020).

16. Jara-Ettinger, J. Theory of mind as inverse reinforcement learning. Curr. Opin. Behav. Sci. 29, 105–110 (2019).

17. Bridgers, S., Jara-Ettinger, J. & Gweon, H. Young children consider the expected utility of others’ learning to decide what to teach. Nat. Hum. Behav. (2019). doi:10.1038/s41562-019-0748-6

18. Liu, S., Ullman, T. D., Tenenbaum, J. B. & Spelke, E. S. Ten-month-old infants infer the value of goals from the costs of actions. Science (80-.). 358, 1038–1041 (2017).

19. Baker, C. L., Saxe, R. & Tenenbaum, J. B. Action understanding as inverse planning. Cognition 113, 329–349 (2009).

20. Baker, C. L., Jara-Ettinger, J., Saxe, R. & Tenenbaum, J. B. Rational quantitative attribution of beliefs, desires and percepts in human mentalizing. Nat. Hum. Behav. 1, 1–10 (2017).

21. Nihonsugi, T., Ihara, a. & Haruno, M. Selective Increase of Intention-Based Economic Decisions by Noninvasive Brain Stimulation to the Dorsolateral Prefrontal Cortex. J. Neurosci. 35, 3412–3419 (2015).

22. Wedekind, C. & Milinski, M. Cooperation through image scoring in humans. Science (80-.). 288, 850–852 (2000).

23. Harbaugh, W. T., Mayr, U. & Burghart, D. R. Neural responses to taxation and voluntary giving reveal motives for charitable donations. Science (80-.). 316, 1622–5 (2007).

24. Niv, Y. et al. Reinforcement learning in multidimensional environments relies on attention mechanisms. J. Neurosci. 35, 8145–8157 (2015).

25. Radulescu, A., Niv, Y. & Ballard, I. Holistic Reinforcement Learning: The Role of Structure and Attention. Trends Cogn. Sci. 23, 278–292 (2019).

26. Leong, Y. C., Radulescu, A., Daniel, R., Dewoskin, V. & Niv, Y. Dynamic Interaction between Reinforcement Learning and Attention in Multidimensional Environments. Neuron 93, 451–463 (2017).

27. Collins, A. G. E. & Frank, M. J. Neural signature of hierarchically structured expectations predicts clustering and transfer of rule sets in reinforcement learning. Cognition 152, 160–169 (2016).

28. Schapiro, A. C., Rogers, T. T., Cordova, N. I., Turk-Browne, N. B. & Botvinick, M. M. Neural representations of events arise from temporal community structure. Nat. Neurosci. 16, 486–492 (2013).

29. Wilson, R. C. & Niv, Y. Inferring relevance in a changing world. Front. Hum. Neurosci. 5, 1–14 (2012).

30. Schapiro, A. C., Turk-Browne, N. B., Norman, K. A. & Botvinick, M. M. Statistical learning of temporal community structure in the hippocampus. Hippocampus 26, 3–8 (2016).

31. Gershman, S. J. & Niv, Y. Learning latent structure: Carving nature at its joints. Curr. Opin. Neurobiol. 20, 251–256 (2010).

32. Kemp, C. & Tenenbaum, J. B. The discovery of structural form. Proc. Natl. Acad. Sci. U. S. A. 105, 10687–10692 (2008).

33. Heckathorn, D. D. The Dynamics and Dilemmas of Collective Action. Am. Sociol. Rev. 61, 250–277 (1996).

34. Axelrod, R. Effective Choice in the Prisoner’ s Dilemma. J. Conflict Resolut. 24, 3–25 (1980).

35. Nash, J. F. Equilibrium Points in n-Person Games. Proc. Natl. Acad. Sci. United States Am. 36, 48–49 (1950).

36. Schwarz, G. Estimating the Dimension of a Model. Ann. Stat. 6, 461–464 (1978).

37. Hampton, A. N., Bossaerts, P. & O’Doherty, J. P. Neural correlates of mentalizing-related computations during strategic interactions in humans. Proc. Natl. Acad. Sci. U. S. A. 105, 6741–6746 (2008).

38. Nosenzo, D., Offerman, T., Sefton, M. & Van Der Veen, A. Discretionary sanctions and rewards in the repeated inspection game. Manage. Sci. 62, 502–517 (2016).

39. FeldmanHall, O. & Shenhav, A. Resolving uncertainty in a social world. Nat. Hum. Behav. 3, (2019).

40. Gershman, S. J., Pouncy, H. T. & Gweon, H. Learning the Structure of Social Influence. Cogn. Sci. 41, 545–575 (2017).

41. Shin, Y. S. & Niv, Y. Biased evaluations emerge from inferring hidden causes. PsyArXiv (2020).

42. Lau, T., Pouncy, H. T., Gershman, S. J. & Cikara, M. Discovering social groups via latent structure learning. J. Exp. Psychol. Gen. 147, 1881–1891 (2018).

43. Gobet, F. et al. Chunking mechanisms in human learning Fernand. Trends Cogn. Sci. 5, (2001).

44. Thalmann, M., Souza, A. S. & Oberauer, K. How does chunking help working memory? J. Exp. Psychol. Learn. Mem. Cogn. 45, 37–55 (2019).

45. Johnson, E. J., Camerer, C., Sen, S. & Rymon, T. Detecting failures of backward induction: Monitoring information search in sequential bargaining. J. Econ. Theory 104, 16–47 (2002).

46. Nagel, R. Unraveling in Guessing Games: An Experimental Study. Am. Econ. Rev. 85, 1313–1326 (1995).

47. Rapoport, A. Individual strategies in a market entry game. Gr. Decis. Negot. 4, 117–133 (1995).

48. Kollock, P. Social Dilemmas: The Anatomy of Cooperation. Annu. Rev. Sociol. 24, 183–214 (1998).

49. Fischbacher, U., Gächter, S. & Fehr, E. Are People Conditionally Cooperative? Evidence from a Public Goods Experiment. Econ. Lett. 71, 397–404 (2001).

50. Kurzban, R. & Houser, D. Experiments investigating cooperative types in humans: A complement to evolutionary theory and simulations. Proc. Natl. Acad. Sci. U. S. A. 102, 1803–1807 (2005).

51. Moore, T. & Zirnsak, M. Neural Mechanisms of Selective Visual Attention. Annu. Rev. Psychol. 68, 47–72 (2017).

52. Caprariello, P. A., Cuddy, A. J. C. & Fiske, S. T. Social structure shapes cultural stereotypes and emotions: A causal test of the stereotype content model. Gr. Process. Intergr. Relations 12, 147–155 (2009).

53. McCauley, C., Stitt, C. L. & Segal, M. Stereotyping: From prejudice to prediction. Psychol. Bull. 87, 195–208 (1980).

54. Raymond S. Nickerson. Confirmation Bias: A Ubiquitous Phenomenon in Many Guises. Rev. Gen. Psychol. 2, 175–220 (1998).

55. Pierson, E. et al. A large-scale analysis of racial disparities in police stops across the United States. Nat. Hum. Behav. (2020). doi:10.1016/j.athoracsur.2014.09.078

56. Gureckis, T. M. et al. psiTurk: An open-source framework for conducting replicable behavioral experiments online. Behav. Res. Methods 48, 829–842 (2016).

57. Licht, A. N. Games Commissions Play : 2×2 Games of International Securities Regulation. Yale J. Int. Law 24, 61–125 (1999).

58. Bramoullé, Y. Anti-coordination and social interactions. Games Econ. Behav. 58, 30–49 (2007).

59. Skyrms, B. The Stag Hunt. Proc. Addresses Am. Philos. Assoc. 75, 31–41 (2001).

60. Fehr, E. & Fischbacher, U. Social norms and human cooperation. Trends Cogn. Sci. 8, 185–190 (2004).

61. Goeree, J. K., Holt, C. A. & Palfrey, T. R. Risk averse behavior in generalized matching pennies games. Games Econ. Behav. 45, 97–113 (2003).

62. Wilson, R. C. & Collins, A. G. Ten simple rules for the computational modeling of behavioral data. Elife 8, (2019).

